# Non-linear Computation Between Neighboring Dendritic Hotspots in Rat CA1 Hippocampal Interneurons

**DOI:** 10.1101/2020.02.06.936989

**Authors:** Attila Kaszás, Zoltán Szadai, Klaudia Spitzer, Gergely Katona, Éva Tóth, Gábor Tamás, Balázs Rózsa

**Affiliations:** Institute of Experimental Medicine of the Hungarian Academy of Sciences, Szigony u. 43, H-1083 Budapest, Hungary; Pázmány Péter Catholic University, The Faculty of Information Technology, Práter u. 50/A, H-1083 Budapest, Hungary; Institut de Neurosciences de la Timone, CNRS UMR 7289, Faculté de Médecine, 27, boulevard Jean Moulin, 13005 Marseille, France; Bioelectronics Department, École des Mines de Saint-Étienne, Campus George Charpak, 880, route de Mimet, 13541 Gardanne cedex, France; Research Group for Cortical Microcircuits of the Hungarian Academy of Sciences, Department of Physiology, Anatomy and Neuroscience, University of Szeged, Közép fasor 52, H-6726 Szeged, Hungary

## Abstract

Dendritic hotspots have been shown to play a determining role as local activity foci during sensory information coding and memory formation in cortical neurons *in vivo*. Here we aim to reveal the characteristics of co-activating more dendritic hotspots themselves within a dynamic time window. Using electrical stimulation and patterned two-photon uncaging for the coincident activation of two neighboring dendritic hotspots in rat hippocampal CA1 interneurons, the evoked local dendritic Ca^2+^ and the associated excitatory postsynaptic potentials (EPSPs) summed supralinearly. Between active hotspots, supralinearity was present in a relatively broad temporal (∼10 ms) and spatial (∼30 µm) window, mediated by NMDA receptors. Though at larger inter-hotspot distances (>30 µm) supralinear summation of EPSPs was replaced by sublinear integration, the summation of dendritic Ca^2+^ responses remained supralinear. The inter-hotspot integration represents a novel signal integration state, extending the interaction distance between dendritic inputs, and allowing long-range coincidence detection.

## INTRODUCTION

Neuronal microcircuits are organized to form cell assemblies that activate dendritic inputs in spatiotemporally clustered patterns (Katona et al., 2011; Kleindienst et al., 2011; Makino and Malinow, 2011; Takahashi et al., 2012). This anatomically structured and temporally organized activity is nonlinearly transformed to action potential output by the dendritic and somatic compartments of downstream neurons (Losonczy and Magee, 2006; Larkum et al., 2009; Katona et al., 2011). During this process, supralinear (Losonczy and Magee, 2006; Larkum et al., 2009; Katona et al., 2011) or sublinear (Abrahamsson et al., 2012; Vervaeke et al., 2012) dendritic integration extends the computational and information storage capacity of neurons according to computational theories (Poirazi et al., 2003). This nonlinear integration is mediated by voltage-gated ion channels and local membrane potential interactions which compartmentalize dendritic arithmetic – initially - into small (∼10 µm) integration subunits (dendritic ‘hotspots’) (Polsky et al., 2004; Katona et al., 2011). If dendrites are bombarded with a higher number of synaptic inputs, these nonlinear interactions can be spatially extended and will generate more global signals, dendritic spikes, which can invade whole dendritic branches (Johnston and Narayanan, 2008; Chiovini et al., 2014). Dendritic hotspots are also formed and stabilized by the compartmentalized biochemical machinery of dendritic plasticity (Chen et al., 2011; Makino and Malinow, 2011; Murakoshi and Yasuda, 2012). Adjacent inputs may encode very different sensory information in pyramidal cells (Jia et al., 2010; Varga et al., 2011; Chen et al., 2013) and interneurons (Chen et al., 2013). Dendritic hotspots may therefore support a new model of associative memory. Whether dendritic hotspots represent independent computational subunits or whether neighboring dendritic hotspots can interact nonlinearly is not known.

In order to support functional compartmentalization, pyramidal neurons provide an anatomical compartment, the dendritic spine (Yuste and Denk, 1995), which is missing in many types of interneurons. However, recent evidence for the functional compartmentalization of Ca^2+^ responses in interneuron dendrites (Goldberg et al., 2003; Rozsa et al., 2004; Camire and Topolnik, 2012; Topolnik and Camire, 2019) has indicated that similar dendritic integration properties may appear locally, as in pyramidal neurons. Correspondingly, sublinear dendritic integration (Abrahamsson et al., 2012; Vervaeke et al., 2012) has been shown in short dendritic segments of interneurons. Moreover, supralinear signal integration between two different input pathways (Calixto et al., 2008), in parvalbumin-positive interneurons (Cornford et al., 2019), in the interneurons of the cerebellum (Tran-Van-Minh et al., 2016), and also in association with dendritic hotspots (Katona et al., 2011; Chiovini et al., 2014), has been demonstrated in hippocampal interneurons.

To gain an insight into neuronal integration states, we investigated a newly discovered intermediate level of dendritic integration which is inserted between the within-hot spot and the axosomatic integration levels, and which could be defined by inter-hot spot interactions in dendrites. Using combined patch-clamp recording, and two-photon imaging and uncaging we have found a strong supralinear summation between the local dendritic and somatic signals of two neighboring dendritic hotspots. This supralinear summation was dominantly mediated by NMDA channels and was replaced by linear and sublinear integration with decreasing spatiotemporal coincidence of hotspot signals.

## RESULTS

### Synaptic integration between dendritic computational subunits

In order to investigate integration between two dendritic hotspots, we first used focal synaptic stimulation. We evoked EPSPs by exciting local axon afferents arriving onto CA1 stratum radiatum interneuron (R/LM-IN) dendrites in two separate dendritic regions (Fig. 1A, see Fig. S1 for cell reconstructions). Two stimulation pipettes were placed juxtaposed to the dendrites (∼ 10 µm away), filled with ACSF, and were followed using infrared scanning gradient contrast microscopy (Wimmer et al., 2004). In contrast to pyramidal neurons, interneurons in the CA1 region of the hippocampus are mostly aspiny (Yuste and Denk, 1995; Goldberg et al., 2003) with a synaptic input density that was approximately 10-fold lower. This resulted in an approximately 100-fold lower success rate in the efficiency of maintaining a stable synaptic stimulation simultaneously in two dendritic regions, as compared to pyramidal cells (Polsky et al., 2004). In order to increase this success rate, stimulation pipettes were repositioned using an automatic, two-photon guided manipulator system until the synaptic stimulation provided stable activation in both dendritic regions (Fig. 1*A*, see Materials and Methods for further details).

**Figure 1.**
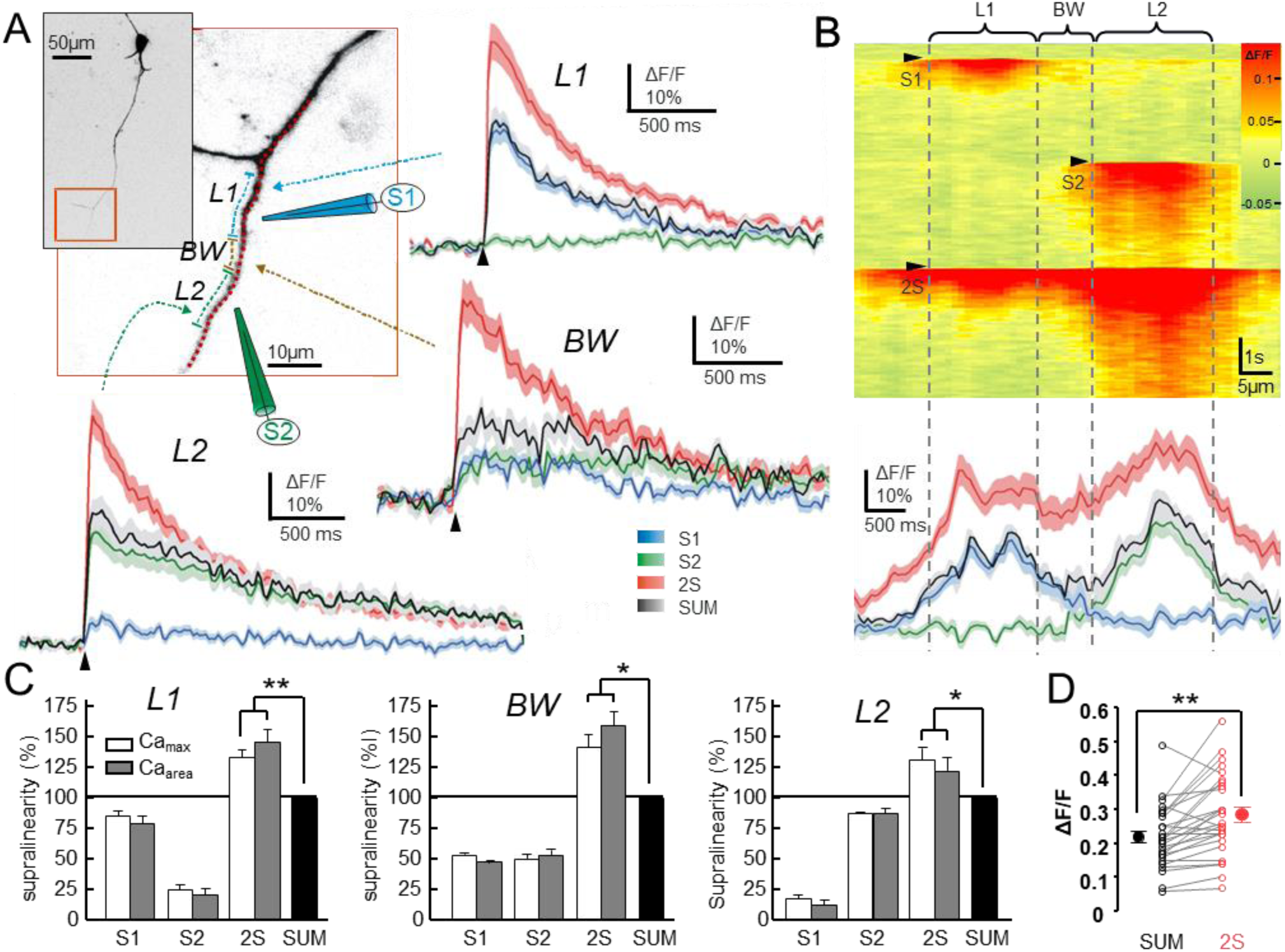
Supralinear integration between neighboring dendritic hotspots. ***A***, Inverted maximum intensity z-stack projection of an R/LM-IN filled with OGB-1 Ca^2+^ indicator. Red box shows the enlarged region of interest with the sketch of the two stimulation pipettes (blue and green). Two dendritic regions (L1 and L2) were activated either separately (defined as S1 and S2 activation, respectively) or simultaneously (defined as 2S) by using focal synaptic stimulation. BW indicates the area between the L1 and L2 regions. Blue, brown, and green dashed lines indicate the L1, BW, and L2 dendritic regions, respectively, where the Ca^2+^ transients were spatially averaged. Average Ca^2+^ transients (mean ± SEM, n=40) for the S1 (blue) and S2 (green) synaptic stimulation in L1, L2, and BW dendritic regions are compared with the simultaneous synaptic stimulation (2S), and with the mathematical sum of the responses for separate S1 and S2 activation (SUM). Red dotted line indicates the path of the free line scan. Black arrowheads indicate time of synaptic stimulation. ***B***, Top, Spatially normalized Ca^2+^ response for the S1, S2, and 2S synaptic stimulations (black triangles) recorded along the red dotted line in A. Bottom, Average spatial distribution of peak Ca^2+^ response (mean ± SEM). ***C***, Average amplitude of Ca^2+^ transient in L1 and L2, and BW (n=7/6 regions/cells), regions for S1, S2 and S2 stimulations compared to the mathematical sum (SUM) of the S1 and S2 stimulation. Transients were normalized to the arithmetic sum in each region. ***D***, Pooled Ca^2+^ amplitudes (mean ± SEM, n=27/9 regions/cells) comparing the measured sum (2S) and arithmetic sum (SUM). Asterisks indicate significance (* p<0.05; ** p<0.001).

The Ca^2+^ signal was acquired using free line scans placed to follow the curvature of the dendrites (Fig. 1*A,B*). Spatially normalized Ca^2+^ signals were plotted as a function of dendritic distance and time, and showed a spatiotemporally compartmentalized activity - dendritic hotspots - following synaptic stimulation. The spatial extension of the hotspots was 9.00 ± 0.87 µm (FWHM, n = 20/8 locations/cells), similar to our previous study performed on CA1 stratum radiatum dendritic innervating interneurons (Katona et al., 2011). In order to understand inter-hot spot interactions, we measured the summation between two neighboring dendritic hotspots for which we selected two dendritic regions (L1 and L2) located on the same, unbranched dendritic segment at a distance of 11.22 ± 2.59 µm (n = 20/8 locations/cells). At this dendritic distance, single synaptic stimulation either at the L1 or L2 location evoked Ca^2+^ transients where the spatial distribution overlapped only partially in the intermediate dendritic region (between the L1 and L2 regions, defined as the BW region, Fig. 1*B*). In order to avoid direct interactions between the stimulus electric fields, the second region was activated with a delay of 0.5 ms during double, near-simultaneous stimulation (2S). At this near-simultaneous synaptic stimulation, the Ca^2+^ responses showed supralinear summation at the L1, L2, and BW regions when compared to the arithmetic sum of Ca^2+^ responses induced by separate activations. Supralinearity was found in the peak amplitude and area of dendritic Ca^2+^ responses in L1 (137.20 ± 6.46% and 151.00 ± 11.39%, respectively; p < 0.001, n = 10/8 regions/cells), and in L2 (121.85 ± 9.85% and 124.38 ± 9.50%, respectively; p = 0.02 and p = 0.01, respectively; n=10/8 regions/cells; Fig. 1). In most cases the two dendritic hotspots were spatially distinct enough to exactly determine supralinearity in the BW region. Interestingly, supralinearity had the highest amplitude - a local maximum - in the BW region (supralinearity for the Ca^2+^ peak amplitude: 148.30 ± 13.46%, for the area: 146.93 ± 13.34%, p = 0.004 and p = 0.01, respectively; n= 5/4 regions/cells; Fig. 1 *B,C*). Simultaneously recorded somatic EPSPs showed supralinear summation with similar amplitude. The summation ratio (SR, see Material and Methods) for the peak and area of the EPSPs was 144.48 ± 19.70 % and 136.05 ± 15.10%, respectively (n = 8 cells, p = 0.02; Fig. 2).

**Figure 2.**
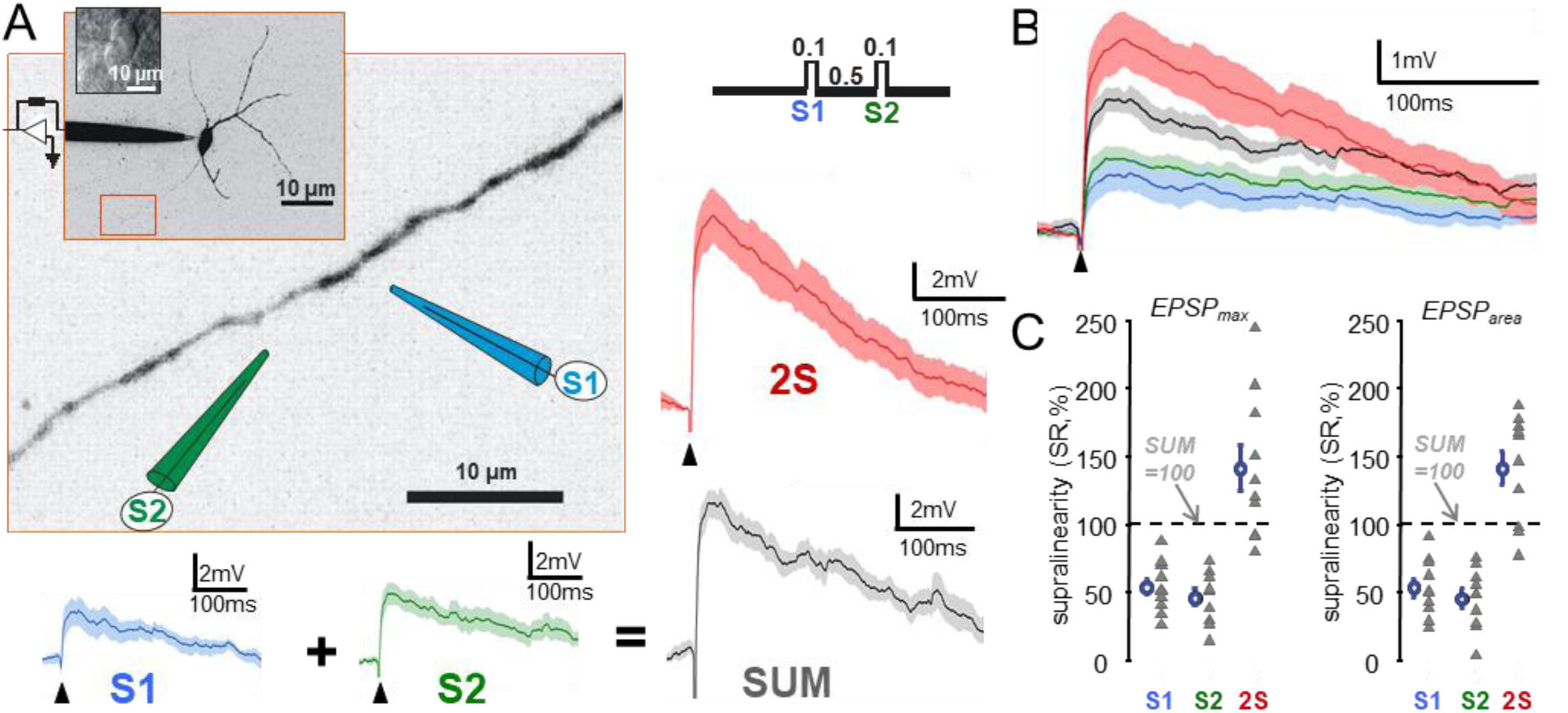
Supralinear integration between neighboring dendritic hotspots is reflected in somatic EPSPs. ***A***, Inverted maximum-intensity z-stack projection of an R/LM-IN with the somatic patch pipette. Red box shows the enlarged dendritic region of interest with the sketch of the two stimulation pipettes (S1, blue; S2, green). Average somatic EPSPs (mean ± SEM) for the separate activation of the first (S1, blue) and second stimulus (S2, green), and the arithmetic sum of the two EPSPs (SUM), are compared to the EPSP of the simultaneous (2S) activation (with 0.5 ms ISI). Inset, transmitted gradient image of the soma was measured with oblique illumination. Top right, stimulation protocol. ***B***, Overlaid transients from *A* showing supralinear summation. ***C***, Summary graph showing average EPSP peak amplitude and area (mean ± SEM) for the recorded and imaged cells (n=9) in percentage of the calculated sum (SUM=100%). Simultaneous synaptic stimulation (2S) showed significant supralinearity (p<0.05). Black arrowhead: time of stimulation.

### Spatially and temporally clustered inputs can reproduce dendritic hotspots and their supralinear interaction

To quantitatively characterize the non-linear interaction between dendritic hotspots, we reproduced two dendritic hotspots by evoking spatially and temporally clustered input patterns using two-photon glutamate uncaging, and compared the mathematical sum of responses induced by separate activation of the two hotspots versus the response of the coincident activation. For this purpose, we wanted to activate the dendritic hotspots by a subthreshold number of unitary inputs. The activation of 20.07 ± 1.03 points (range 8 to 30 points, n = 45/14 locations/cells; Fig. 3*A*, see Materials and Methods) resulted in somatically subthreshold EPSPs. The spatial distribution of the uncaging-evoked dendritic hotspots was not significantly different from that of the focal electric stimulation-evoked hotspots (synaptic stimulation: 9.00 ± 0.87 µm FWHM, uncaging-evoked: 9.79 ± 0.59 µm FWHM, p = 0.95, n = 20/8 regions/cells, and n = 45/14 regions/cells, respectively), and also well reproduced the spontaneously emerged hotspots that we detected in our previous study of R/LM-INs (Katona et al., 2011). One of the advantages of the patterned two-photon activation over focal synaptic stimulation is that contamination of local signals by the stimulation of axons running to the unobserved parts of dendrites can be fully excluded. Moreover, the use of two-photon uncaging eliminates the potential contribution of presynaptic mechanisms. When two dendritic hotspots placed 15.59 ± 0.77 µm apart (range: 9.03 to 29.47 µm, n = 56/14 regions/cells) were near-synchronously stimulated (1 ms inter-stimulus interval), Ca^2+^ responses recorded along the dendrites were supralinear in both hotspots (L1 and L2) as compared to the arithmetic sum of the responses induced by separate activation of the two hotspots (supralinearity was 120.37 ± 4.31% and 123.36 ± 4.81% in the peak amplitude and the area of the Ca^2+^, respectively; n = 56/14 regions/cells; p < 0.001; Fig. 3). Similarly to the data obtained with synaptic stimulation, the supralinearity had a local maximum in the intermediate dendritic region (BW region), where supralinearity in peak amplitude and area was 122.38 ± 5.65% and 134.16 ± 9.81%, respectively (n=18/10 regions/cells; p_amplitude_ < 0.001, p_area_ = 0.001 Fig. 3). Uncaging-evoked EPSPs (uEPSPs) recorded simultaneously at the soma also showed significant supralinearity in their summation ratio. The SR of the peak amplitude and the area of uEPSPs was 110.08 ± 4.16% and 125.88 ± 7.25%, respectively (n = 14 cells; p_area_ < 0.001, p_amplitude_ = 0.006). These data agree well with the synaptic stimulation data, further confirming that neighboring computational subunits, dendritic hotspots, can interact nonlinearly. Hence hotspots provide one more intermediate state in dendritic integration before the signal reaches the axosomatic integration domain.

**Figure 3.**
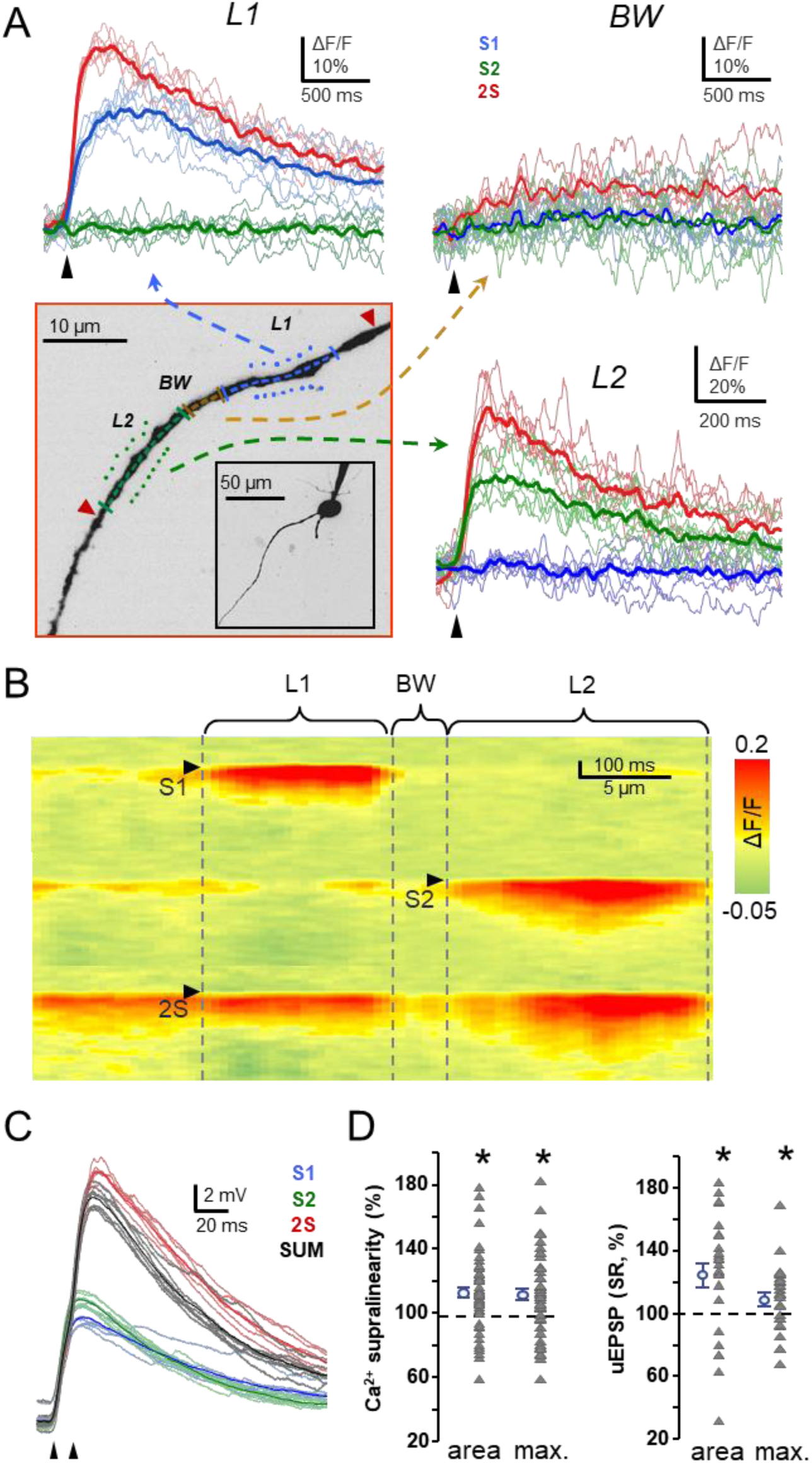
Spatio-temporally clustered excitatory input packages can reproduce supralinear integration between dendritic hotspots. ***A***, Bottom left, inverted maximum-intensity z-stack projection of an R/LM-IN dendrite showing the two clusters of inputs used for patterned glutamate activation in L1 (blue cluster) and L2 (green cluster) dendritic regions. Red triangles indicate the two end points of the dendritic segment, measured with free-line scanning. Dashed arrows are pointing to individual (thin traces) and average (thick traces) uncaging-evoked Ca^2+^ transients which were induced by either separate (S1 and S2, blue and green traces, respectively) or simultaneous (2S, red traces) activation of the two dendritic input clusters. Transients were measured in the L1, L2, and BW dendritic regions as indicated. ***B***, Average Ca^2+^ response recorded between the two red triangles along the dendritic segment shown in *A* following either separate (S1 and S2) or simultaneous (2S) activation of the two dendritic input clusters shown in *A*. ***C***, Simultaneously recorded individual (thin) and mean (thick) somatic uEPSPs for the separate (S1: blue, S2: green) and for the simultaneous (2S: red) activation of the two clusters in *A*. SUM: calculated sum (black). ***D***, Left, summary graph showing the mean supralinearity (2S/(S1+S2)*100) in summation of the peak amplitude and area of Ca^2+^ responses (n=52/12 hotspots/cells). Right, summation ratio (2S-S1)/S2*100) of uEPSP peaks and areas (n=12 cells). Double stimulation induced significant supralinearity in both the Ca^2+^ EPSP responses (asterisk: p<0.05). Black arrowheads indicate time of uncaging.

Next, we investigated the spatial extent of the interaction. The supralinearity of EPSP areas decreased rapidly as a function of dendritic distance between the two input clusters, following an exponential decay pattern (Fig. 4*E-F*). Moreover, sublinearity replaced supralinearity at dendritic distances over 50 µm. Interestingly, the fall in the supralinearity of Ca^2+^ responses followed a more modest tendency, and never crossed the level of linearity.

**Figure 4.**
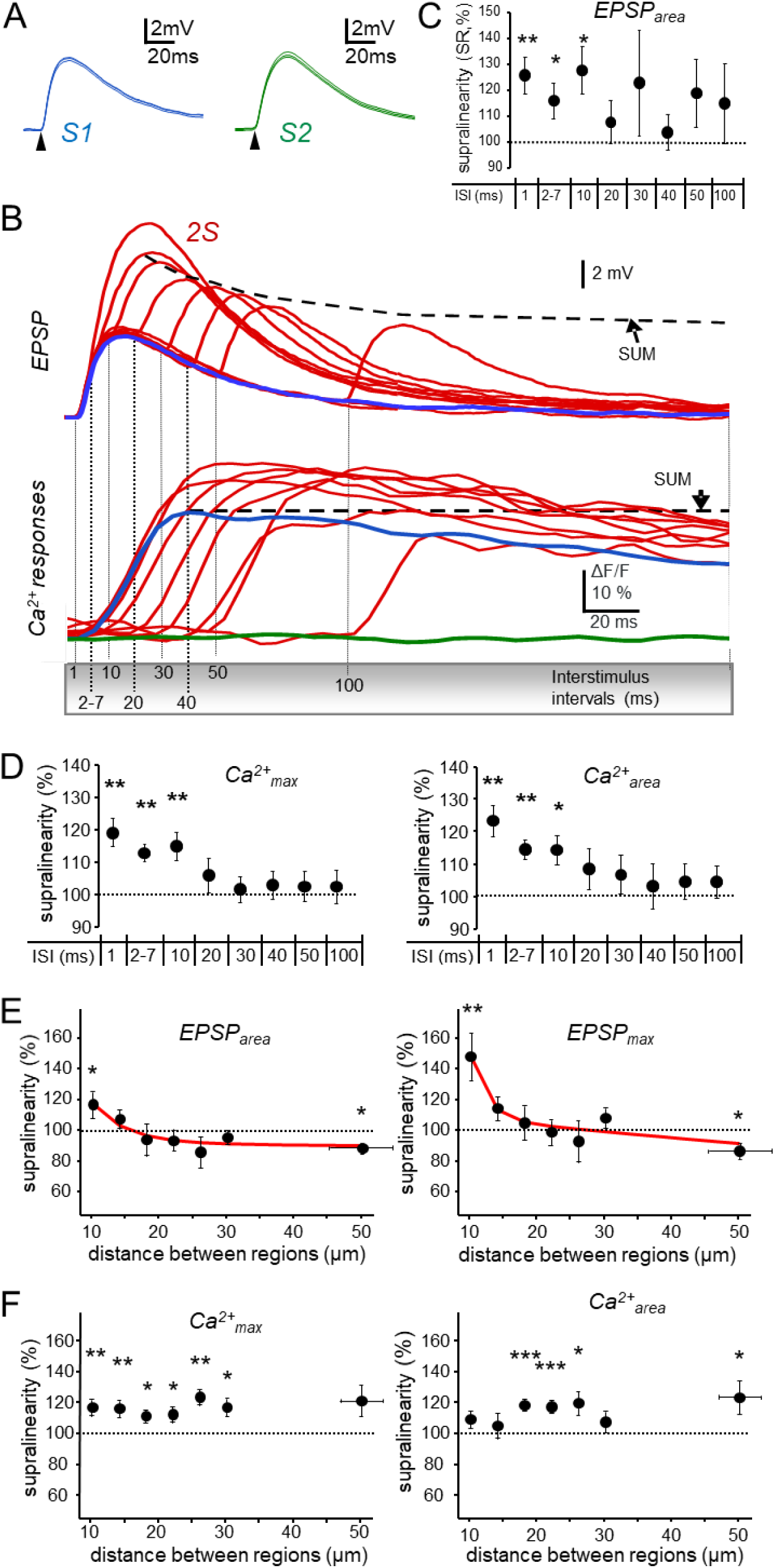
Supralinear hot spot integration depends on temporal and spatial synchronicity of inputs. ***A***, Representative uEPSPs evoked with the first (S1) or second (S2) input cluster, similarly to Fig. 3, in two neighboring dendritic hot spot regions. Black arrowheads indicate uncaging time. ***B***, The two neighboring hotspots were activated with different ISI and the summed response of the double activation (2S: red) was compared to the individual responses of separate hot spot activation (S1: blue, S2: green). Representative traces are given for the uEPSPs (top) and for the simultaneously recorded dendritic Ca^2+^ signals (bottom) measured in the first dendritic hot spot region. Dashed lines indicate peak of the arithmetic sums (SUM). ***C***, Statistics for the timing-dependent supralinearity of uEPSP areas. Summation ratio shows supralinearity for 1-10 ms ISI. ***D***, Mean supralinearity in summation of the peak and area of Ca^2+^ responses as a function of ISI. See also Table 1. ***E***, Summation ratio of EPSP areas (left) and amplitudes (right) as a function of distance between the two input clusters. Red line shows exponential fit. ***F***, The same as in *E* but for the ratio of the supralinearity in summation of the peak amplitude and area of Ca^2+^ responses. Note the preserved supralinearity as a function of distance. Dataset was binned and the last 5/10 data points were pooled on *E* and *F*. Asterisks indicate significance (* p<0.05, ** p<0.01).

To investigate the temporal dependence of inter hot spot interactions we used the same uncaging protocol, but varied the temporal delay between the activation of the two hotspots by between 0.5 and 100 ms (Fig. 4). The somatic uEPSPs showed supralinearity for synchronous activation of the two hotspots in a 10 ms temporal window (Fig. 4*B,C*). Furthermore, the simultaneously recorded Ca^2+^ responses showed a similar supralinear increase in their amplitude and area (Fig. 4*D*, Table 1). This relatively long integration window is on the scale of the life-time of dendritic hotspots emerging spontaneously (Takahashi et al., 2012) and indicates that NMDARs mediate these mechanisms (Schiller et al., 2000; Katona et al., 2011).

**Table 1.**
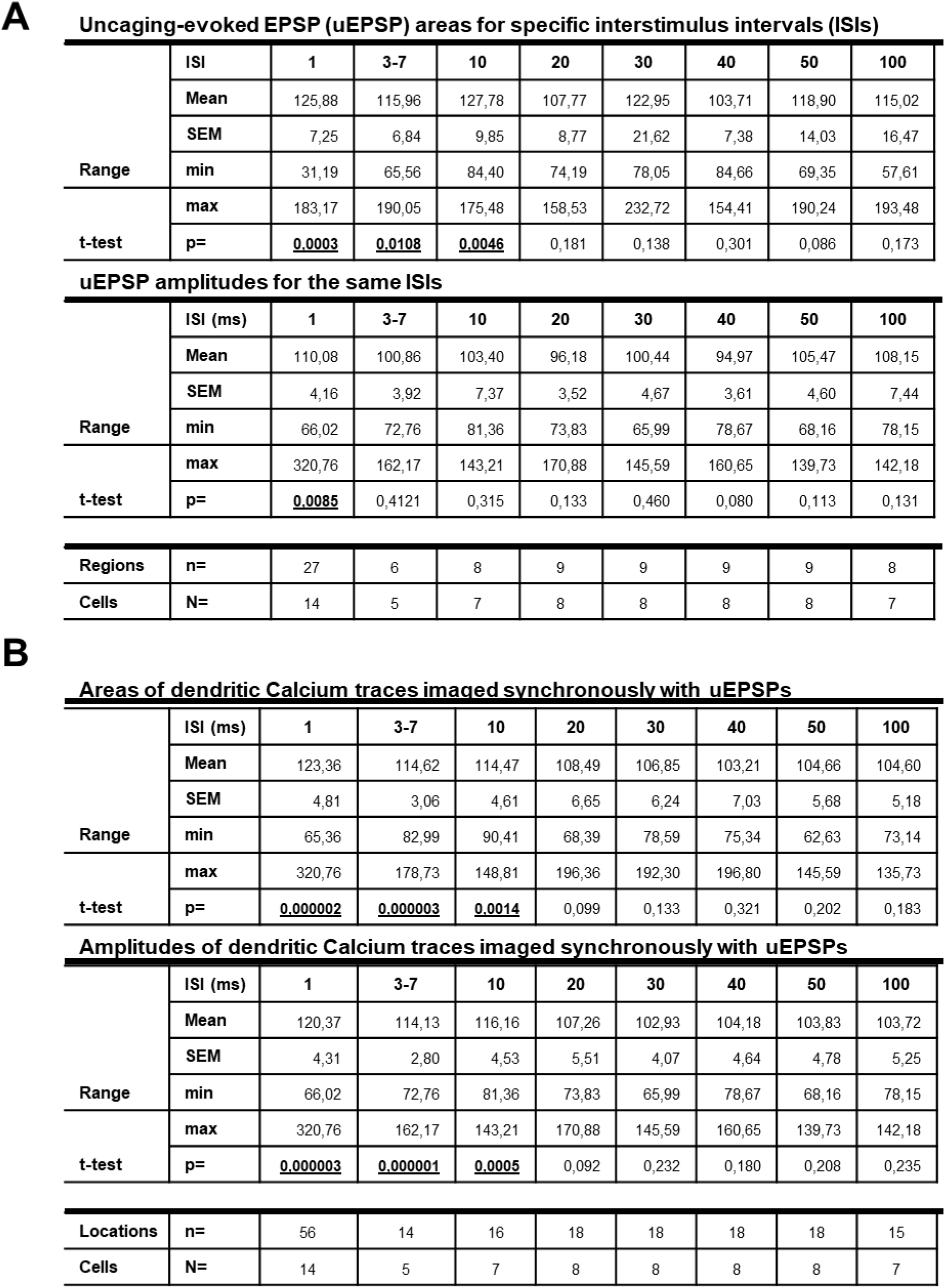
Signal integration of inputs with different intersimulus intervals. Tables show values for the summation of two uncaging-evoked events separated by a defined intersimulus interval (ISI). A) Somatically recorded uEPSP areas and maxima. Values are in %Summation Ratio [%SR=(2S-S1)/S2*100]. B) Local Calcium trace areas and maxima for the different (ISIs). Values are in %control [%control=2S/(S1+S2)*100].

### NMDARs shape inter hotspot integration

The large dendritic Ca^2+^ signals, the elongated EPSPs, and the relatively long temporal window of inter-hotspots interactions suggested that NMDARs were involved in the mechanism of the non-linear inter-hot spot integration (Schiller et al., 2000; Katona et al., 2011). Earlier works have confirmed that NMDARs have a role in dendritic hot spot generation (Katona et al., 2011; Kleindienst et al., 2011), we therefore applied the NMDAR blocker AP5, which decreased the uEPSP amplitude and area of individual hot spot responses by 27.21 ± 9.06% and 40.67 ± 7.81%, respectively (Fig. 5*C*, Fig. S5, n = 10/4 regions/cells, p < 0.05). Similarly, both the amplitude and the area of the simultaneously recorded dendritic Ca^2+^ responses decreased by 51.07 ± 10.14% and 70.82 ± 7.32% (Fig. 5*A*, n = 10/4 regions/cells, p < 0.05). Then, we also examined the change in the supralinear interaction of two dendritic hotspots. We observed that the summation ratio of uEPSP amplitudes and area decreased by 14.79 ± 5.88% and 20.79 ± 5.93% from linear summation, respectively (n = 5/4 regions/cells, p < 0.05) in the presence of AP5, and this decrease was also reflected in the simultaneously recorded dendritic Ca^2+^ responses, where supralinearity turned to sublinearity in summation of amplitude and area, decreasing by 9.08 ± 9.66% and 21.45 ± 6.51% from linear summation, respectively (n = 5/4 regions/cells, p < 0.05).

**Figure 5.**
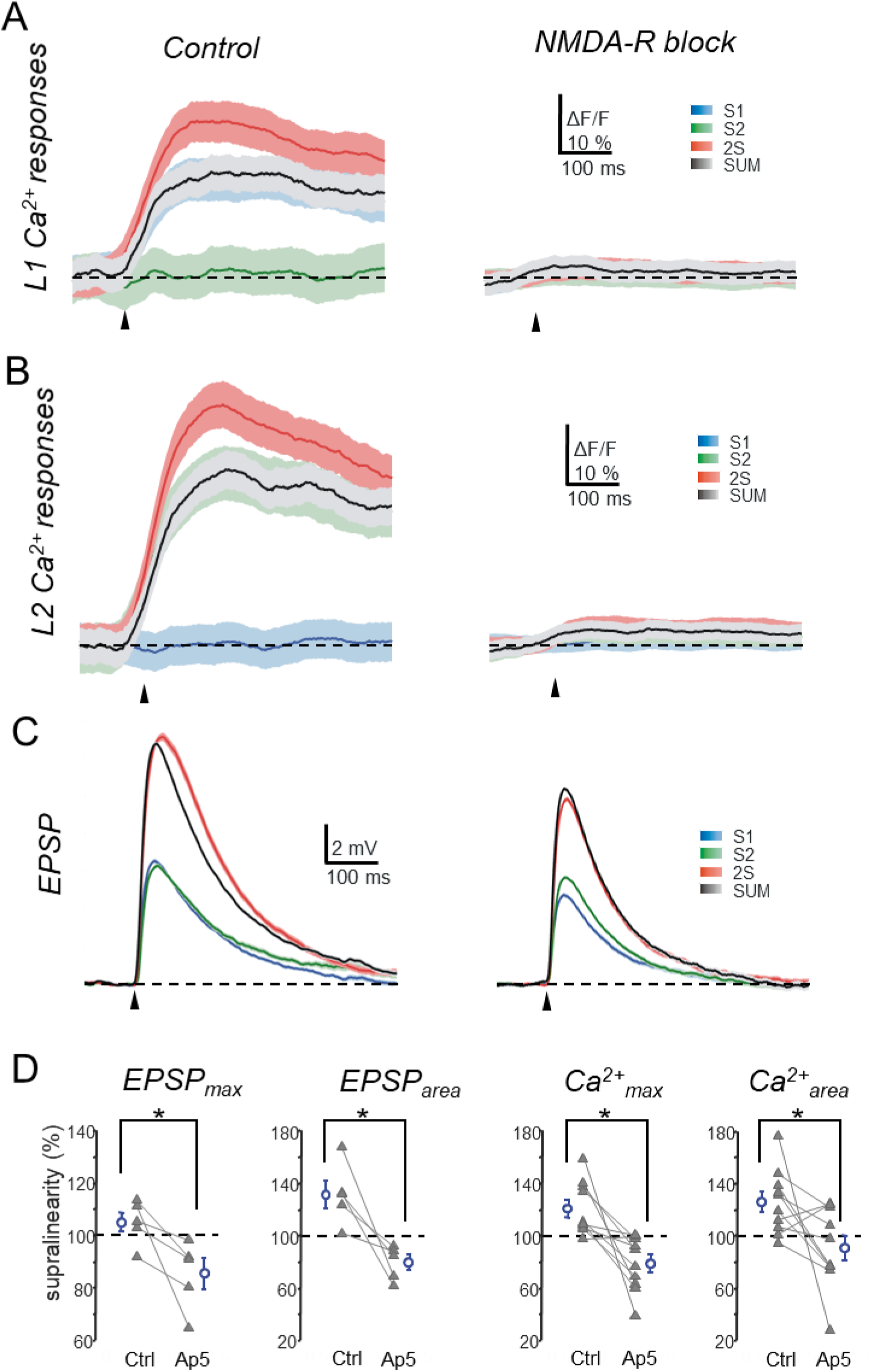
Inter hot spot interaction is mediated by NMDARs. ***A, B***, Average dendritic Ca^2+^ transients from the two stimulated regions (mean ± SEM) in the presence of NMDAR blocker AP5 (control: left, Ap5: right). Responses for the first (S1, blue) and second stimulus (S2, green), their arithmetic sums (SUM, black line) and the activity after double stimulation (2S, red line) are shown. Black arrowheads indicate the time of the stimulation. ***C***, Same as in *A*, and B, but for the simultaneously recorded somatic uEPSPs. ***D***, Summary of the effect of NMDAR blockage. The graphs show the change in summation ratio for uEPSP amplitude (1st) and area (2nd), and for dendritic Ca^2+^ transient amplitudes (3rd) and area (4th) in the presence of AP5 (n=5/4 regions/cells). Asterisks indicate significance (*p<0.05, paired t-test).

## DISCUSSION

Our conclusions are based on a series of functional experiments performed using whole-cell patch clamping, two-photon imaging, and two-photon uncaging of the caged-glutamate compound, DNI-glutamate•TFA (Chiovini et al., 2014; Palfi et al., 2018). First, we found that the somatically recorded EPSPs evoked by focal synaptic stimulation of local synaptic afferents arriving onto two discrete, but neighboring, dendritic regions show supralinear summation in R/LM-INs. Furthermore, the locality of signals was confirmed in two-photon imaging of Ca^2+^ responses, where the synchronous synaptic stimulation of these inputs induced a local, supralinear Ca^2+^ influx at the two activated hot spot regions, and also in the neighboring dendritic segments. Consistent with the results of the focal synaptic stimulation, when two separate clusters of glutamatergic inputs were near simultaneously activated, we observed supralinear summation of both the somatic EPSPs and the local Ca^2+^ signals. Interestingly, changing the relative distance between the input clusters revealed that there is a long-range (>50 µm) interaction between the local dendritic Ca^2+^ responses, while summation of somatic EPSPs turned rapidly from supralinear to sublinear as a function of distance. Moreover, the supralinearity of responses was apparent only in a time window of 1 to 10 ms. Finally, our pharmacological experiments revealed a determining role of NMDARs in the inter-hot spot interaction.

A growing body of evidence supports the conclusion that neuronal networks are wired in a way that means that temporally synchronized cell assemblies can activate dendritic hotspots in pyramidal cells (Kleindienst et al., 2011; Makino and Malinow, 2011; Takahashi et al., 2012) and interneurons (Katona et al., 2011; Camire and Topolnik, 2014; Tran-Van-Minh et al., 2016). The observed synchronization of inputs can affect local events on two different timescales. Firstly, local membrane potential depolarization can spread rapidly and activate voltage-dependent ion channels such as the voltage-dependent Na^+^, Ca^2+^, or K^+^ channels (Losonczy and Magee, 2006; Larkum et al., 2009), or NMDARs (Schiller et al., 2000; Katona et al., 2011). This local interaction can result in supralinear or sublinear summation of the neighboring synaptic inputs (Schiller et al., 2000; Losonczy and Magee, 2006; Larkum et al., 2009; Katona et al., 2011; Vervaeke et al., 2012; Chiovini et al., 2014; Topolnik and Camire, 2019). Secondly, the interaction between synchronized inputs can bloom into medium- or long-term events by affecting the local concentration of second messenger molecules (Harvey et al., 2008; Fitzpatrick et al., 2009), thus leading to the generation of spatially restricted long-term plasticity (LTP), (Camire and Topolnik, 2014). Because neighboring presynaptic inputs represent very different sensory information (Jia et al., 2010; Chen et al., 2011; Chadderton et al., 2014), the compartmentalized plasticity can attract axons (Chklovskii et al., 2004) and connect cells representing variable sensory information into an assembly. Therefore, dendritic hotspots appear to be local organizer elements essential to associative memory. The spatial extension of dendritic hotspots in interneurons (Goldberg et al., 2003; Katona et al., 2011; Chen et al., 2013) and in pyramidal cells (Jia et al., 2010; Kleindienst et al., 2011; Takahashi et al., 2012) has been shown to be small, being on the scale of a few micrometers, also with small spatial variance; however, the spillover of the neurotransmitters (Hires et al., 2008), and the local spread of the diffusible signaling molecules (Fitzpatrick et al., 2009) and dendritic membrane potential (Krueppel et al., 2011) can be more extended, allowing long-range interactions between neighboring computational subunits. Indeed, we found a significant nonlinear interaction in the summation of somatic EPSPs during coincident activation of two dendritic hotspots in a range of about 30 µm. In the first ∼ 20 µm range, nonlinearity was dominated by supralinearity, which was then replaced by linear and sublinear summation over 50 µm. Similar sublinear integration between low-density excitatory inputs has been shown previously in cortical interneurons (Tamas et al., 2002), and it may reflect shunting between neighboring conductances. In the first 20 µm range, the shunting effect of neighboring conductances is complemented by the nonlinear effect of voltage-gated channels, which reverse the direction of summation toward supralinearity, according to our measurements. In contrast to somatic EPSPs, the supralinearity in summation of local dendritic Ca^2+^ responses did not decline as a function of dendritic distance between two hotspots, in fact it had a maximum between two hotspots. This mismatch in the nonlinearity of summation between somatic EPSPs and dendritic Ca^2+^ responses can be explained by the role of dendritic NMDARs. The local dendritic Ca^2+^ signals were dominated by highly nonlinear NMDARs, but NMDARs contributed much less to the somatic EPSPs. A similar mismatch in the amplitude of nonlinearity was found in other neurons during the summation of somatic EPSPs and local dendritic Ca^2+^ responses (Makara and Magee, 2013).

Regarding the temporal scale of the integration, the supralinearity of responses was apparent in a temporal window about 10 ms long. Temporal windows of this length are characteristic for coincidence detection (Calixto et al., 2008) in the neural circuitry. Previous data indicated that interneurons are able to take a substantial role in controlling oscillatory activity of the hippocampus (Klausberger, 2009). Therefore, this short coincidence window may allow them to integrate input assemblies within a single oscillation cycle during different types of brain oscillations, such as gamma or theta oscillations.

All previous work has indicated that pyramidal neurons can express supralinear integration in their dendrites (Stuart, 1999; Schiller et al., 2000; Losonczy and Magee, 2006; Larkum et al., 2009; Branco and Hausser, 2011), but interneurons paint a more colorful picture according to recent data, because their dendritic integration can be both supra- and sublinear. For example, cerebellar Golgi cells (Vervaeke et al., 2012) and molecular layer interneurons (Tran-Van-Minh et al., 2016), stellate cells (Abrahamsson et al., 2012) and cortical interneurons (Tamas et al., 2002) have sublinear integration. In contrast, interneuronal integration can also be supralinear when the number of activated inputs reaches a given threshold (Cornford et al., 2019). This happens, for example, when the physiological network is better preserved and generates a higher background input activity (Katona et al., 2011; Chiovini et al., 2014), or when multiple paths are integrated simultaneously (Jarsky et al., 2005; Calixto et al., 2008). Our experiments have revealed supralinear signal integration in R/LM-IN dendrites, with a previously unexplored dendritic integration state which is inserted between the layers of dendritic and axo-somatic integrator domains. We propose that this third level of dendritic integration extends the currently accepted two-layer neuronal integration model (Poirazi et al., 2003; Larkum et al., 2009), because neighboring hotspots can interact with each other on a much longer spatial scale before the information reaches the axo-somatic integration domain, following nonlinear integration within individual dendritic hotspots, in a second non-linear step within individual dendritic hotspots. As neighboring dendritic inputs can represent very different sensory information in pyramidal neurons (Jia et al., 2010) and GABAergic interneurons (Chen et al., 2013), this novel integration state can connect more distal dendritic inputs and in this way can spatially extend the functional basis of associative memory in dendrites.

## MATERIALS AND METHODS

### Slice preparation and electrophysiology

Acute hippocampal slices were prepared from 16-24-day-old Wistar rats using isoflurane anesthesia followed by swift decapitation, in accordance with the Hungarian Act of Animal Care and Experimentation (1998; XXVIII, section 243/1998.). Coronal (300 µm) brain slices were cut with a Vibratome 3000 (Vibratome) and stored at room temperature in artificial cerebrospinal fluid (ACSF) (in mM: 126 NaCl, 2.5 KCl, 2 CaCl_2_, 2 MgCl_2_, 1.25 NaH_2_PO_4_, 26 NaHCO_3_, and 10 glucose) as previously described (Rozsa et al., 2004).

Hippocampal neurons in the CA1 stratum radiatum near the border of the stratum lacunosum-moleculare (R/LM-IN) were visualized using 850 nm infrared lateral illumination. Current-clamp recordings were made at 32 °C (MultiClamp 700B, Digidata 1440; Molecular Devices) with glass electrodes (6-10 MΩ) filled with (in mM: 125 K-gluconate, 20 KCl, 10 HEPES, 10 phosphocreatine di(tris)-salt, 0.3 Na-GTP, 4 Mg-ATP, 10 NaCl, 0.06 Oregon Green BAPTA-1). 5 mg/ml biocytin was added to the pipette solution for later morphological identification of the recorded cells. Cells with a resting membrane potential of below −50 mV were accepted. The recorded cells were classified as regular spiking, Schaffer collateral-associated cells and apical dendrite innervating interneurons according to their electrophysiological properties (resting membrane potential V_m_ = −68.43±1.6 mV, firing adaptation 53.6±3.17%) (Lamsa et al., 2007), two-photon image stacks, and 3D light microscopic reconstructions (Fig. S1) (Sik et al., 1995; Freund and Buzsaki, 1996; Ascoli et al., 2008). The NMDA-receptor (NMDAR) antagonist D, L-AP5 (60 µM) was applied to the bath. Unless otherwise stated, all chemicals and drugs were purchased from Sigma. Data acquisition was performed using either pClamp8 or pClamp10 (Molecular Devices), and MES (Femtonics Ltd.) software.

### Multiphoton imaging

Imaging was performed using a two-photon laser scanning system (Femto2D-uncage, Femtonics Ltd., Budapest, Hungary). The imaging lasers (Mai Tai HP Deep See, SpectraPhysics, or Chameleon Ultra II, Coherent) was set to 830 nm for the focal electrical stimulation, and to 840 nm in the uncaging experiments. Free line scans were placed following the curvature of long dendritic segments to monitor the Ca^2+^ signals elicited by focal electric stimulation (elicited by two electrodes, located at different relative distances). The scanning time was less than 6 ms, which allowed us to analyze the spatiotemporal properties of Ca^2+^ compartments along long dendritic segments. Regions of interest were scanned at a constant speed, while intermediate sections were jumped over within ∼60 µs (Multiple Line-Scan mode of scanning, Femtonics Ltd.), resulting in a higher signal-to-noise ratio than obtained by a raster scan of the same region of interest. Imaging of Ca^2+^ traces started at 20-30 min after break-in to allow for proper filling of the cells with the dye OGB-1. At the end of each experiment, a series of images across the depth of the volume encompassing the imaged neuron, another stack of the imaged dendrite and a low-magnification infrared image of the hippocampus were taken to determine the localization of the neuron and inputs. To monitor patch-clamp conditions and to check for potential photo damage during imaging experiments, we induced a burst of five action potentials (200-400 pA, 5 ms; APs were evoked at 35 Hz) and monitored the amplitude of the somatic APs, and also the local dendritic Ca^2+^ signals, induced by backpropagating action potentials (bAPs).

### Synaptic stimulation

For the synaptic stimulation as a first step we calibrated the three axes of the coordinate system of the patch-clamp micromanipulators (Luigs & Neumann) by holding the stimulation electrodes against the three axes of the coordinate system of the two-photon microscope in a MATLAB-based program. Following the initial manual marking of the tip of the pipettes in the two-photon images to calibrate the origin of the two coordinate systems, the system automatically moved the pipettes at a given speed to the preselected target locations where synaptic stimulation was performed. Within the tissue, the software allowed only movements parallel to the pipette axis: this minimized tissue damage. The target locations for synaptic stimulation were selected according to the two-photon image of the dendrite. The whole process was monitored using the overlaid transmitted infrared image of the pipette and the two-photon fluorescence images of the region of interest.

Focal synaptic stimulation was performed as described earlier (Rozsa et al., 2004). Briefly, 6-9 MΩ glass electrodes filled with ACSF were placed at a distance of 5-25 µm from the dendrite (stimulation: 0.1 ms, 10-50 V, 0.5 ms pulse interval at double pulses; Supertech Ltd.). The GABA_A_ receptor blocker bicuculline (20 µM) added to the bath when using focal electrical stimulation. All evoked excitatory postsynaptic potentials (EPSPs) were verified for synaptic delay.

### Calculation of supralinear summation of somatic EPSPs

For the somatic recordings, we calculated the summation ratio (SR) in percentage using the formula

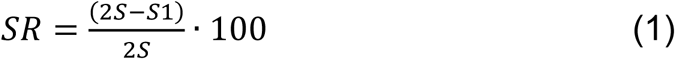

Here, *2S* denotes the amplitude (or area) of the somatic EPSP following stimulation in both dendritic hot spot regions at a certain interstimulus interval (ISI= 0.5; 1; 3-7;10; 20; 30; 40; 50; 100 ms), while *S1* and *S2* are the amplitude (or area) of the somatic EPSP after the first and second stimulus, respectively. Measurement of *S1, S2* and *2S* responses, and thereby the corresponding synaptic stimulations, were separated by 2 s in order to avoid temporal overlaps between these responses (Figure 1B).

In order to calculate nonlinearity in Ca^2+^ responses, we first, defined the border of the two activated dendritic hot spot regions (L1 and L2, Figure 1A and B) at the half maxima of the peak amplitude responses induced by separate stimulation (*S1* and *S2*). The nonlinearity in Ca^2+^ signal summation was then calculated as:

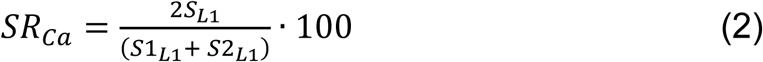

Here, *S1*_*L1*_ and *S2*_*L1*_ denote the responses at location L1 after the first and second stimulus, respectively, and *2S*_*L1*_ denotes the response in region L1 following near-simultaneous activation in both regions with a given inter-stimulus interval. The same calculation was performed in the L2 region, and also between the L2 and L1 regions (“BW” region). The arithmetic sum of the peak amplitude of any two Ca^2+^ responses can differ from the peak amplitude of the arithmetic sum of the same Ca^2+^responses (therefore, for example, the sum of the S1 and S2 bars in Figure 1C is also not quite 100%).

### Multiphoton glutamate uncaging

After achieving whole-cell mode and filling R/LM-INs with 60 µM OGB-1, the bath solution was exchanged to an ACSF containing 4-methoxy-5,7-dinitroindolinyl (DNI)-caged L-glutamate trifluoroacetate (DNI-glutamate•TFA; 2.5 mM) (Chiovini et al., 2014; Palfi et al., 2018), or 4-methoxy-7-nitroindolinyl (MNI)-caged L-glutamate (MNI-glutamate; 2.5 mM; Tocris), or 4-methoxy-7-nitroindolinyl (MNI)-caged L-glutamate trifluoroacetate (MNI-glutamate•TFA; 2.5 mM; Femtonics Ltd.). Photolysis of caged glutamate was performed with 720 nm (for MNI-glutamates) or 740 nm (for DNI-glutamate) ultrafast, pulsed laser light (Mai Tai HP Deep See, SpectraPhysics, or Chameleon, Coherent). The intensity of the uncaging laser beam was controlled with an electro-optical modulator built in the Femto2D-uncage microscope. Chromatic aberration was compensated for at the focal plane, and the incoming uncaging laser beam was aligned using two motorized mirrors in order to overlap imaging and uncaging point spread functions (PSFs) with 100 nm radial and 300 nm axial precision. Two-photon imaging was interleaved with glutamate uncaging periods when the scanner jumped to the maximum 30 selected locations (within 60 µs jump time) and returned back to the scanning trajectory thereafter. The position of the uncaging sites was finely adjusted in order to keep the relative distance between uncaging locations and the dendritic segment stable. The distance of uncaging locations from the dendrite was also monitored by measuring the green fluorescence level during the photostimulation. Locations with a fluorescence artifact during uncaging over a given critical level (∼2000 arbitrary units at 90% PMT voltage saturation) were moved or removed to decrease dendritic phototoxicity. Stimulus artifacts caused by the uncaging in fluorescence curves or spatially normalized images were removed *post hoc*. Photolysis of caged glutamate was performed in clustered patterns (0.69 ± 0.04 µm between inputs) along the dendrite. Amplitude of high osmolar ACSF-induced unitary EPSPs was 0.55 ± 0.12 mV (n = 4). Uncaging clusters consisting of 20.07 ± 1.03 points (range 8-30 points, n = 43/14 regions/cells), which were used to evoke somatically subthreshold EPSPs, were equivalent with 12.37 ± 0.95 unitary inputs (range 8-31 inputs, n=43/14 regions/cells).

### Data analysis and statistics

Unless otherwise indicated, data are reported as mean ± SEM. To test for significance, we used the Student’s t-test at a significance level of p≤0.05.

## FIGURE LEGENDS

**Supplementary Figure S1.**
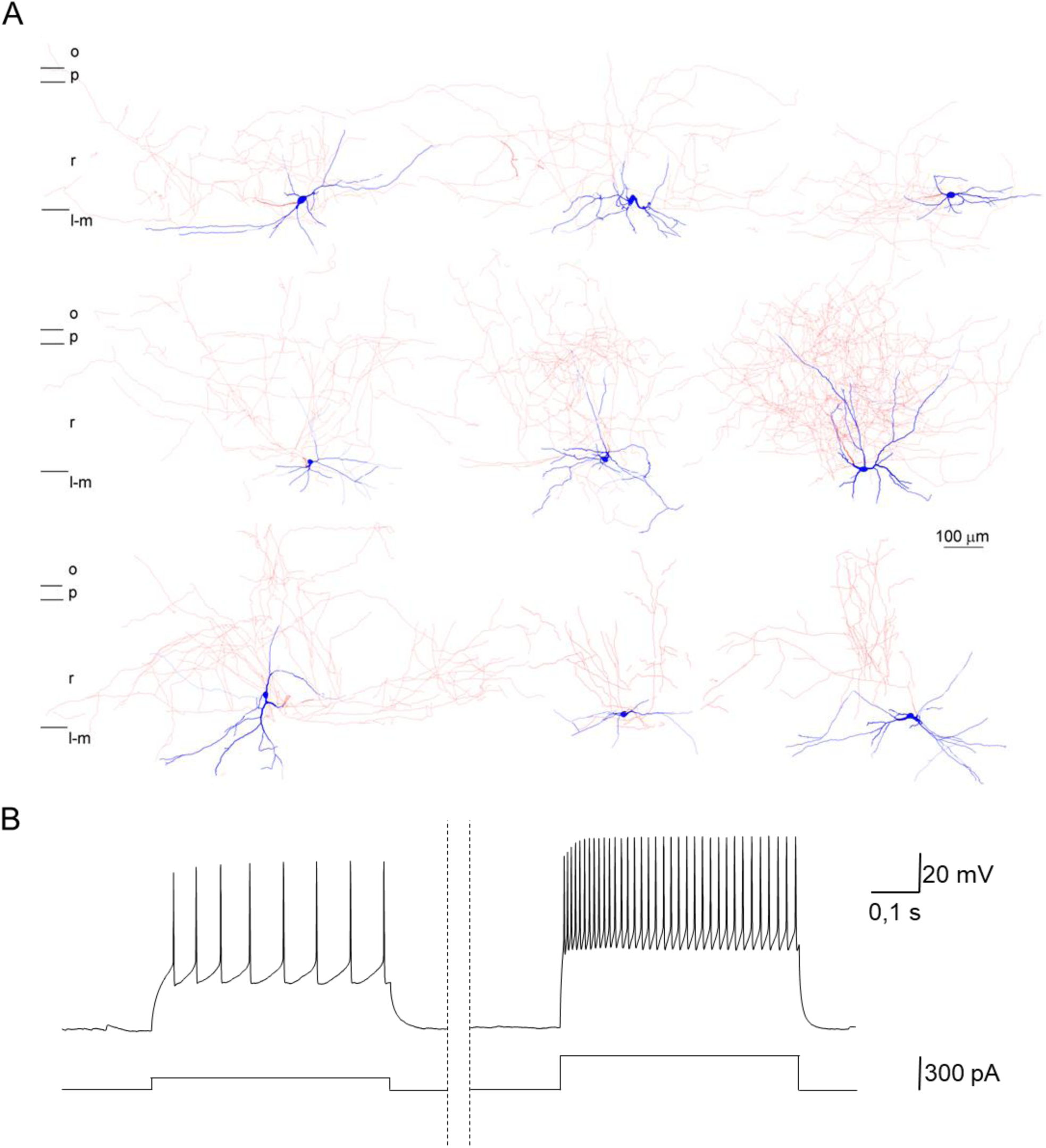
Anatomical and electrophysiological characterization of the recorded cells. **A**, Anatomical reconstruction of the biocytin-filled interneurons showing their position in the hippocampus along with their axonal and dendritic arborization. **B**, Ramp test protocol for electrophysiological characterization of the recorded cells with threshold (*left panel*) and maximum firing frequency (*right panel*) stimuli.

## ADDITIONAL INFORMATION

## Author contributions

A.K. and Z.Sz. contributed equally, B.R. and A.K. designed research; A.K., K.S., Z.Sz., G.T. and É.T. performed research; A.K., K.S., Z.Sz., and B.R. analyzed data; G.K. and B.R. constructed the imaging setup; A.K. and B.R. wrote the paper.

## REFERENCES

Abrahamsson, T., L. Cathala, K. Matsui, R. Shigemoto and D. A. Digregorio (2012). Thin dendrites of cerebellar interneurons confer sublinear synaptic integration and a gradient of short-term plasticity. Neuron 73(6): 1159–1172. DOI: 10.1016/j.neuron.2012.01.027

Ascoli, G. A., L. Alonso-Nanclares, S. A. Anderson, G. Barrionuevo, R. Benavides-Piccione, A. Burkhalter, G. Buzsaki, B. Cauli, J. Defelipe, A. Fairen, D. Feldmeyer, G. Fishell, Y. Fregnac, T. F. Freund, D. Gardner, E. P. Gardner, J. H. Goldberg, M. Helmstaedter, S. Hestrin, F. Karube, Z. F. Kisvarday, B. Lambolez, D. A. Lewis, O. Marin, H. Markram, A. Munoz, A. Packer, C. C. Petersen, K. S. Rockland, J. Rossier, B. Rudy, P. Somogyi, J. F. Staiger, G. Tamas, A. M. Thomson, M. Toledo-Rodriguez, Y. Wang, D. C. West and R. Yuste (2008). Petilla terminology: nomenclature of features of GABAergic interneurons of the cerebral cortex. Nat Rev Neurosci 9(7): 557–568.

Branco, T. and M. Hausser (2011). Synaptic integration gradients in single cortical pyramidal cell dendrites. Neuron 69(5): 885–892. DOI: 10.1016/j.neuron.2011.02.006

Calixto, E., E. J. Galvan, J. P. Card and G. Barrionuevo (2008). Coincidence detection of convergent perforant path and mossy fibre inputs by CA3 interneurons. J Physiol 586(Pt 11): 2695–2712.

Camire, O. and L. Topolnik (2012). Functional compartmentalisation and regulation of postsynaptic Ca2+ transients in inhibitory interneurons. Cell Calcium 52(5): 339–346. DOI: 10.1016/j.ceca.2012.05.001

Camire, O. and L. Topolnik (2014). Dendritic calcium nonlinearities switch the direction of synaptic plasticity in fast-spiking interneurons. J Neurosci 34(11): 3864–3877. DOI: 10.1523/JNEUROSCI.2253-13.2014

Chadderton, P., A. T. Schaefer, S. R. Williams and T. W. Margrie (2014). Sensory-evoked synaptic integration in cerebellar and cerebral cortical neurons. Nat Rev Neurosci 15(2): 71–83. DOI: 10.1038/nrn3648

Chen, T. W., T. J. Wardill, Y. Sun, S. R. Pulver, S. L. Renninger, A. Baohan, E. R. Schreiter, R. A. Kerr, M. B. Orger, V. Jayaraman, L. L. Looger, K. Svoboda and D. S. Kim (2013). Ultrasensitive fluorescent proteins for imaging neuronal activity. Nature 499(7458): 295–300. DOI: 10.1038/nature12354

Chen, X., U. Leischner, N. L. Rochefort, I. Nelken and A. Konnerth (2011). Functional mapping of single spines in cortical neurons in vivo. Nature 475(7357): 501–505. DOI: 10.1038/nature10193

Chen, X., C. D. Nelson, X. Li, C. A. Winters, R. Azzam, A. A. Sousa, R. D. Leapman, H. Gainer, M. Sheng and T. S. Reese (2011). PSD-95 is required to sustain the molecular organization of the postsynaptic density. J Neurosci 31(17): 6329–6338. DOI: 10.1523/JNEUROSCI.5968-10.2011

Chiovini, B., G. F. Turi, G. Katona, A. Kaszas, D. Palfi, P. Maak, G. Szalay, M. F. Szabo, G. Szabo, Z. Szadai, S. Kali and B. Rozsa (2014). Dendritic spikes induce ripples in parvalbumin interneurons during hippocampal sharp waves. Neuron 82(4): 908–924. DOI: 10.1016/j.neuron.2014.04.004

Chklovskii, D. B., B. W. Mel and K. Svoboda (2004). Cortical rewiring and information storage. Nature 431(7010): 782–788. DOI: nature03012 [pii] 10.1038/nature03012

Cornford, J. H., M. S. Mercier, M. Leite, V. Magloire, M. Hausser and D. M. Kullmann (2019). Dendritic NMDA receptors in parvalbumin neurons enable strong and stable neuronal assemblies. Elife 8. DOI: 10.7554/eLife.49872

Fitzpatrick, J. S., A. M. Hagenston, D. N. Hertle, K. E. Gipson, L. Bertetto-D’Angelo and M. F. Yeckel (2009). Inositol-1,4,5-trisphosphate receptor-mediated Ca2+ waves in pyramidal neuron dendrites propagate through hot spots and cold spots. J Physiol 587(Pt 7): 1439–1459. DOI: jphysiol.2009.168930 [pii] 10.1113/jphysiol.2009.168930

Freund, T. F. and G. Buzsaki (1996). Interneurons of the hippocampus. Hippocampus 6(4): 347–470.

Goldberg, J. H., G. Tamas, D. Aronov and R. Yuste (2003). Calcium microdomains in aspiny dendrites. Neuron 40(4): 807–821.

Harvey, C. D., R. Yasuda, H. Zhong and K. Svoboda (2008). The spread of Ras activity triggered by activation of a single dendritic spine. Science 321(5885): 136–140. DOI: 1159675 [pii] 10.1126/science.1159675

Hires, S. A., Y. Zhu and R. Y. Tsien (2008). Optical measurement of synaptic glutamate spillover and reuptake by linker optimized glutamate-sensitive fluorescent reporters. Proc Natl Acad Sci U S A 105(11): 4411–4416. DOI: 0712008105 [pii] 10.1073/pnas.0712008105

Jarsky, T., A. Roxin, W. L. Kath and N. Spruston (2005). Conditional dendritic spike propagation following distal synaptic activation of hippocampal CA1 pyramidal neurons. Nat Neurosci 8(12): 1667–1676. DOI: nn1599 [pii] 10.1038/nn1599

Jia, H., N. L. Rochefort, X. Chen and A. Konnerth (2010). Dendritic organization of sensory input to cortical neurons in vivo. Nature 464(7293): 1307–1312. DOI: 10.1038/nature08947

Johnston, D. and R. Narayanan (2008). Active dendrites: colorful wings of the mysterious butterflies. Trends Neurosci 31(6): 309–316. DOI: 10.1016/j.tins.2008.03.004

Katona, G., A. Kaszas, G. F. Turi, N. Hajos, G. Tamas, E. S. Vizi and B. Rozsa (2011). Roller Coaster Scanning reveals spontaneous triggering of dendritic spikes in CA1 interneurons. Proc Natl Acad Sci U S A 108(5): 2148–2153. DOI: 10.1073/pnas.1009270108

Klausberger, T. (2009). GABAergic interneurons targeting dendrites of pyramidal cells in the CA1 area of the hippocampus. Eur J Neurosci 30(6): 947–957.

Kleindienst, T., J. Winnubst, C. Roth-Alpermann, T. Bonhoeffer and C. Lohmann (2011). Activity-dependent clustering of functional synaptic inputs on developing hippocampal dendrites. Neuron 72(6): 1012–1024. DOI: 10.1016/j.neuron.2011.10.015

Krueppel, R., S. Remy and H. Beck (2011). Dendritic integration in hippocampal dentate granule cells. Neuron 71(3): 512–528. DOI: S0896-6273(11)00500-9 [pii] 10.1016/j.neuron.2011.05.043

Lamsa, K. P., J. H. Heeroma, P. Somogyi, D. A. Rusakov and D. M. Kullmann (2007). Anti-Hebbian long-term potentiation in the hippocampal feedback inhibitory circuit. Science 315(5816): 1262–1266.

Larkum, M. E., T. Nevian, M. Sandler, A. Polsky and J. Schiller (2009). Synaptic integration in tuft dendrites of layer 5 pyramidal neurons: a new unifying principle. Science 325(5941): 756–760. DOI: 10.1126/science.1171958

Losonczy, A. and J. C. Magee (2006). Integrative properties of radial oblique dendrites in hippocampal CA1 pyramidal neurons. Neuron 50(2): 291–307.

Makara, J. K. and J. C. Magee (2013). Variable dendritic integration in hippocampal CA3 pyramidal neurons. Neuron 80(6): 1438–1450. DOI: 10.1016/j.neuron.2013.10.033

Makino, H. and R. Malinow (2011). Compartmentalized versus global synaptic plasticity on dendrites controlled by experience. Neuron 72(6): 1001–1011. DOI: 10.1016/j.neuron.2011.09.036

Murakoshi, H. and R. Yasuda (2012). Postsynaptic signaling during plasticity of dendritic spines. Trends Neurosci 35(2): 135–143. DOI: 10.1016/j.tins.2011.12.002

Palfi, D., B. Chiovini, G. Szalay, A. Kaszas, G. F. Turi, G. Katona, P. Abranyi-Balogh, M. Szori, A. Potor, O. Frigyesi, C. Lukacsne Haveland, Z. Szadai, M. Madarasz, A. Vasanits-Zsigrai, I. Molnar-Perl, B. Viskolcz, I. G. Csizmadia, Z. Mucsi and B. Rozsa (2018). High efficiency two-photon uncaging coupled by the correction of spontaneous hydrolysis. Org Biomol Chem 16(11): 1958–1970. DOI: 10.1039/c8ob00025e

Poirazi, P., T. Brannon and B. W. Mel (2003). Pyramidal neuron as two-layer neural network. Neuron 37(6): 989–999.

Polsky, A., B. W. Mel and J. Schiller (2004). Computational subunits in thin dendrites of pyramidal cells. Nat Neurosci 7(6): 621–627. DOI: 10.1038/nn1253 nn1253 [pii]

Rozsa, B., T. Zelles, E. S. Vizi and B. Lendvai (2004). Distance-dependent scaling of calcium transients evoked by backpropagating spikes and synaptic activity in dendrites of hippocampal interneurons. J Neurosci 24(3): 661–670.

Schiller, J., G. Major, H. J. Koester and Y. Schiller (2000). NMDA spikes in basal dendrites of cortical pyramidal neurons. Nature 404(6775): 285–289. DOI: 10.1038/35005094

Sik, A., M. Penttonen, A. Ylinen and G. Buzsaki (1995). Hippocampal CA1 interneurons: an in vivo intracellular labeling study. J Neurosci 15(10): 6651–6665.

Stuart, G. S. N.; Hausser, M. (1999). Dendrites, Oxford University Press.

Takahashi, N., K. Kitamura, N. Matsuo, M. Mayford, M. Kano, N. Matsuki and Y. Ikegaya (2012). Locally synchronized synaptic inputs. Science 335(6066): 353–356. DOI: 10.1126/science.1210362

Tamas, G., J. Szabadics and P. Somogyi (2002). Cell type- and subcellular position-dependent summation of unitary postsynaptic potentials in neocortical neurons. J Neurosci 22(3): 740–747.

Topolnik, L. and O. Camire (2019). Non-linear calcium signalling and synaptic plasticity in interneurons. Curr Opin Neurobiol 54: 98–103. DOI: 10.1016/j.conb.2018.09.006

Tran-Van-Minh, A., T. Abrahamsson, L. Cathala and D. A. DiGregorio (2016). Differential Dendritic Integration of Synaptic Potentials and Calcium in Cerebellar Interneurons. Neuron 91(4): 837–850. DOI: 10.1016/j.neuron.2016.07.029

Varga, Z., H. Jia, B. Sakmann and A. Konnerth (2011). Dendritic coding of multiple sensory inputs in single cortical neurons in vivo. Proc Natl Acad Sci U S A 108(37): 15420–15425. DOI: 10.1073/pnas.1112355108

Vervaeke, K., A. Lorincz, Z. Nusser and R. A. Silver (2012). Gap junctions compensate for sublinear dendritic integration in an inhibitory network. Science 335(6076): 1624–1628. DOI: 10.1126/science.1215101

Wimmer, V. C., T. Nevian and T. Kuner (2004). Targeted in vivo expression of proteins in the calyx of Held. Pflugers Arch 449(3): 319–333. DOI: 10.1007/s00424-004-1327-9

Yuste, R. and W. Denk (1995). Dendritic spines as basic functional units of neuronal integration. Nature 375(6533): 682–684.

